# Targeted Modification of the Antimicrobial Peptide DGL13K Reveals a Naturally Optimized Sequence for Topical Applications

**DOI:** 10.1101/2025.09.16.676538

**Authors:** Sven-Ulrik Gorr

**Affiliations:** Department of Diagnostic and Biological Sciences, University of Minnesota School of Dentistry, Minneapolis Minnesota, 55455, U.S.A

**Keywords:** Allo-isoleucine, Antimicrobial peptide, Bacteria, DGL13K, Hemolysis, Myristoylation, Peptide modification, Peptide sequence, Polyethylene glycol

## Abstract

Antimicrobial peptides are potential alternatives to conventional antibiotics, primarily due broad spectrum activity and low propensity for inducing bacterial resistance. However, their clinical translation faces challenges, including peptide stability and potential mammalian cell toxicity. This study centers on DGL13K, an all D-amino acid peptide, which overcomes proteolytic susceptibility and demonstrates notable stability and broad-spectrum bactericidal activity without inducing de novo bacterial resistance.

This work aimed to enhance the therapeutic properties of DGL13K by using targeted modifications to increase antimicrobial potency and decrease toxicity, as determined by hemolysis. DGL13K derivatives were synthesized and tested, involving amino acid substitutions, stereochemical alterations, and N-terminal functionalization with polyethylene glycol (PEG) or myristoylate. While some modifications altered bacterial specificity and reduced hemolytic activity, none of the tested alterations resulted in a substantial overall improvement compared to the parent DGL13K sequence. Furthermore, the antibacterial efficacy of DGL13K and its variants was significantly inhibited in the presence of 50% serum, suggesting limitations for systemic applications.

The findings suggest that the DGL13K sequence, derived from an evolutionarily selected protein, is already highly optimized. Given its stability, broad-spectrum efficacy, in vivo activity, low resistance profile, and high safety margin, DGL13K is a promising therapeutic candidate for topical/localized infections.

## INTRODUCTION

### Antimicrobial peptides

Antimicrobial peptides (AMPs) are a diverse group of host-defense molecules that are found throughout nature in organisms from bacteria and fungi to plants and animals, serving as an ancient component of innate immunity [1-3]. Combined with synthetic versions of natural peptides and de novo designed peptides, this class of antimicrobial molecules displays a wide variety of sequences, structures and antimicrobial properties [4-9]. AMPs are prominent in mucosal surfaces where they serve as a first line of defense against invading microorganisms. As an example, we have cataloged at least 45 distinct antimicrobial peptides and proteins in the oral cavity [10,11]. Interestingly, this rich environment of AMPs, allows the colonization by commensal organisms while invading microbes are effectively killed in this environment. Thus, it was already noted in the 1930’s that saliva allows the growth of oral bacteria while killing non-oral bacteria [12]. These properties suggest that human AMPs could be exploited as a new class of antimicrobial therapeutics with low potential for pathogen resistance while triggering fewer side-effects by limiting the disruption of the commensal host microbiome.

Despite their promise as an alternative to traditional antibiotics, a number of challenges have been identified in the attempt to develop AMPs for clinical use [13-15]. As an example, natural peptides are susceptible to proteolytic degradation although this can be largely overcome by the use of unnatural and D-amino acids in synthetic peptides [16-19]. Many AMPs target the bacterial cell membrane [20,21] but, due to similarities with mammalian cell membranes, this is not always a specific target and mammalian cell toxicity has been cited as a concern for clinical development [22-24].

### Design of novel therapeutic AMPs

Thousands of AMPs have been identified and can be found in several online databases [7-9,25]. These peptides typically contain hydrophobic and cationic amino acids but no consensus sequence has been identified for antibacterial activity and the roles of intervening amino acids is poorly understood. Nevertheless, these databases can be queried for common themes that define AMPs [26]. In addition, recent machine learning and broader AI models have been developed to better predict novel AMP sequences [27-30], although the ultimate success of these approaches remains to be demonstrated [31].

### Design of DGL13K

Our laboratory has developed the second-generation AMP DGL13K, which was derived from the anti-inflammatory salivary protein BPIFA2 [32-34]. BPIFA2 is a Leu-rich protein [35] that is abundant in rodent [36,37] and dog [38] salivary glands/saliva and present in human saliva [39], albeit in relatively low amounts [40]. This protein belongs to a family of antibacterial and anti-inflammatory proteins that are found in multiple mucosal surfaces and secretions [41-43]. We noted that BPIFA2 facilitates bacterial aggregation and LPS binding [44-46]. To identify the active domain of BPIFA2, the protein was compared to active domains in the related proteins BPI and LBP, which also exhibit LPS-binding activity [32,47]. The initial peptide, GK7 was extended to develop GL13NH2 that shows anti-LPS and bacteria agglutinating activity [33,44,46].

Similar to BPIFA2, GL13NH2 causes bacterial agglutination, which is able to prevent the spread of *Pseudomonas* infection in a plant model [33]. However, GL13NH2 does not kill the bacteria. To achieve bactericidal activity, the positive charge of the peptide was increased by substituting three polar amino acids with Lys residues [34]. The resulting peptide, GL13K (now named LGL13K) exhibits bactericidal activity against most Gram negative bacteria but is relatively inactive against Gram positive bacteria [19,34,48-51].

LGL13K is susceptible to bacterial proteases, which led to the design of the stereo-isomer DGL13K [16]. DGL13K resists proteolytic degradation [16,19] and has antibacterial activity against all tested strains of both Gram negative and Gram positive bacteria, including *A. baumanii* (6 strains) [49], *Enterococcus faecalis* (7 strains) [19], *K. pneumoniae* (7 strains) [49], *Porphyromonas gingivalis* (2 strains) [50], *P. aeruginosa* (9 strains) [16,48,49], *S. aureus* (2 strains) [48], and *Streptococcus gordonii* (3 strains) [19]. Recent unpublished data also show efficacy against *Bacteroides fragilis, Clostridioides difficile, Enterobacter cloacae, Enterococcus faecium* and *Escherichia coli*, thereby completing the ESKAPEE pathogens [52]. In addition, DGL13K shows activity against drug-resistant Gram-negative bacteria [49], including extended-spectrum beta-lactamase (ESBL)-producing and carbapenemase (KPC)-producing *K. pneumoniae*, multi-drug resistant and extensively drug-resistant *P. aeruginosa*, and extensively drug-resistant *A. baumannii* carrying metallo-beta-lactamases. Activity against drug-resistant Gram-positive bacteria includes methicillin-resistant *S. aureus* (MRSA) [48] and vancomycin-resistant *E. faecalis* (VRE) [19].

### Structure-function studies of GL13K

LDGL13K is predominantly found in a random coil conformation in the absence of membranes, the peptide adopts an α-helical structure from residue K5 to K11 in the presence of dodecylphosphocholine micelles [53]. In the presence of negatively charged lipid bilayers, the peptide is predominantly found in a ß-sheet structure [53,54]. These ß-sheets can assemble into nanofibrils, which may be the active form of the peptide [55,56]. Rather than forming membrane pores, the relatively short peptide (13 amino acids) disrupts the structure of the bacterial membrane by removing lipid micelles [53,54].

### Resistance

No tested bacteria have developed resistance to DGL13K: Prolonged treatment with sub-inhibitory concentrations (0.5xMIC) of DGL13K does not lead to resistance in *P. aeruginosa* [49], *S. aureus* [51], *E. faecalis* or *S. gordonii* [19]. Remarkably, when *S. aureus* is treated with the L- or D-isomer of GL13K, they gain resistance to LGL13K but not DGL13K, which remains effective against the selected bacteria [51].

The goal of this study was to use targeted modifications of the peptide sequence and modification by functional groups to test if the activity of the peptide can be increased while reducing toxicity to mammalian cells.

## Material and Methods

### Bacteria

*Pseudomonas aeruginosa* Xen41 and *Staphylococcus aureus* Xen36 were obtained from Xenogen (Alameda, CA; now Revvity, Waltham, MA) and Revvity, respectively and stored at −80°C in 10% glycerol. *P. aeruginosa* were cultured in Luria-Bertani broth while *S. aureus* were cultured in Todd-Hewitt Broth (Difco, Franklin Lakes, NJ) overnight at 37°C and shaking at 200 rpm. Cultures typically reached an optical density at 600 nm of 1.7 for *P. aeruginosa* and 1.3 for *S. aureus*.

### Peptide sequences

Synthetic peptides and N-terminally modified peptides were purchased from Bachem (Torrance, CA) or Aaptec (Louisville, KY) (**Table 1**). Peptide identity and purity was verified by the manufacturer by mass spectrometry and RP-HPLC, respectively. Unless otherwise noted, the peptides were C-terminally amidated and delivered as a TFA salt at >95% purity. Peptide stock solutions were prepared at 10 mg/ml in dH_2_O or 0.01% acetic acid and stored at 4°C until use. We have recently reported that the stock solutions are stable for at least 2 years under these conditions [51].

### Heat stability

Peptides were diluted to 1 mg/ml in sterile 10%PBS (1 part PBS, 9 parts dH2O) and heated for 60 min in a 0.5 ml microcentrifuge tube in a water bath set at 80°C. Control samples were similarly incubated at room temperature. The samples were used for MIC assays, as described below.

### Minimal Inhibitory Concentration

MIC assays were performed as previously described [51]. Briefly, overnight cultures of *P. aeruginosa* Xen41 were diluted to 10^5^ CFU/ml in Mueller-Hinton Broth while *S. aureus* Xen36 were similarly diluted in Todd-Hewitt Broth. Bacteria (100 µl) were added to 20 µl of 2-fold peptide dilutions in 10% PBS in 96-well polypropylene culture plates. The plates were incubated overnight at 37°C with gentle rocking and then the optical density at 630 nm (OD630) was recorded in a BioTek Synergy HT plate reader (BioTek, Winooski, VT; now Agilent, Santa Clara, CA). Bioluminescence was recorded for quality control of the bioluminescent bacteria.

In some MIC assays, the culture medium was supplemented with 60% heat-inactivated fetal calf serum (final assay concentration: 50% serum).

### Hemolysis

Lysis of human red blood cells (Innovative Research; Novi, MI) was determined as previously described [48].

**Table 1:**
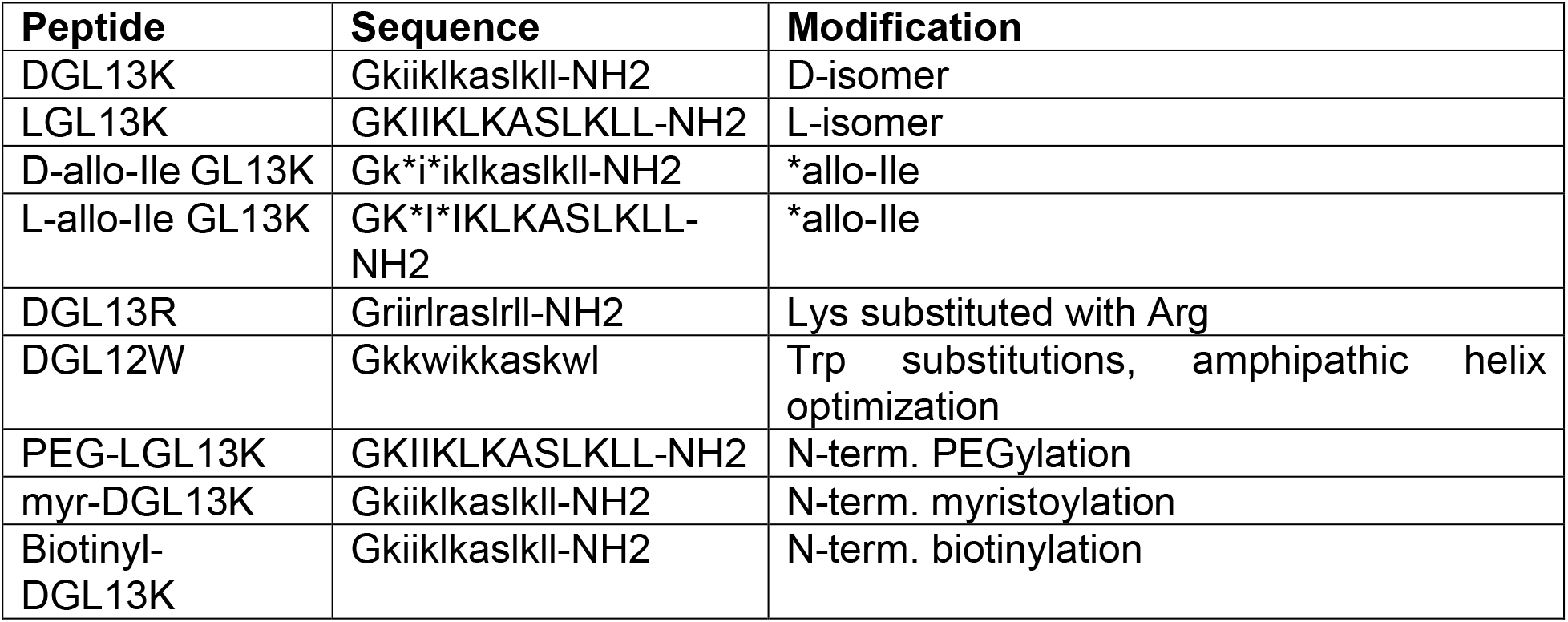
Peptide modifications. Table 1 lists the peptides discussed in this paper. Sequences are shown as upper case letters for L-amino acids and lower case for D-amino acids. Allo-Ile is marked by *. -NH2 – C-terminal amidation. PEG – polyethylene glycol; myr - myristoyl.

## RESULTS AND DISCUSSION

### Stability of DGL13K in aqueous solution

Peptides are typically considered inherently unstable in aqueous solutions [57]. However, an aqueous solution of DGL13K did not lose antibacterial activity after storage at 4°C for more than 2 years [51]. Similarly, there is no loss of activity when a solution of DGL13K is heated at 80°C for an hour (**Fig. 1**). Together these results point to the robustness of DGL13K, which overcomes the frequent concern that AMPs are not sufficiently stable for therapeutic use [58].

**Fig 1.**
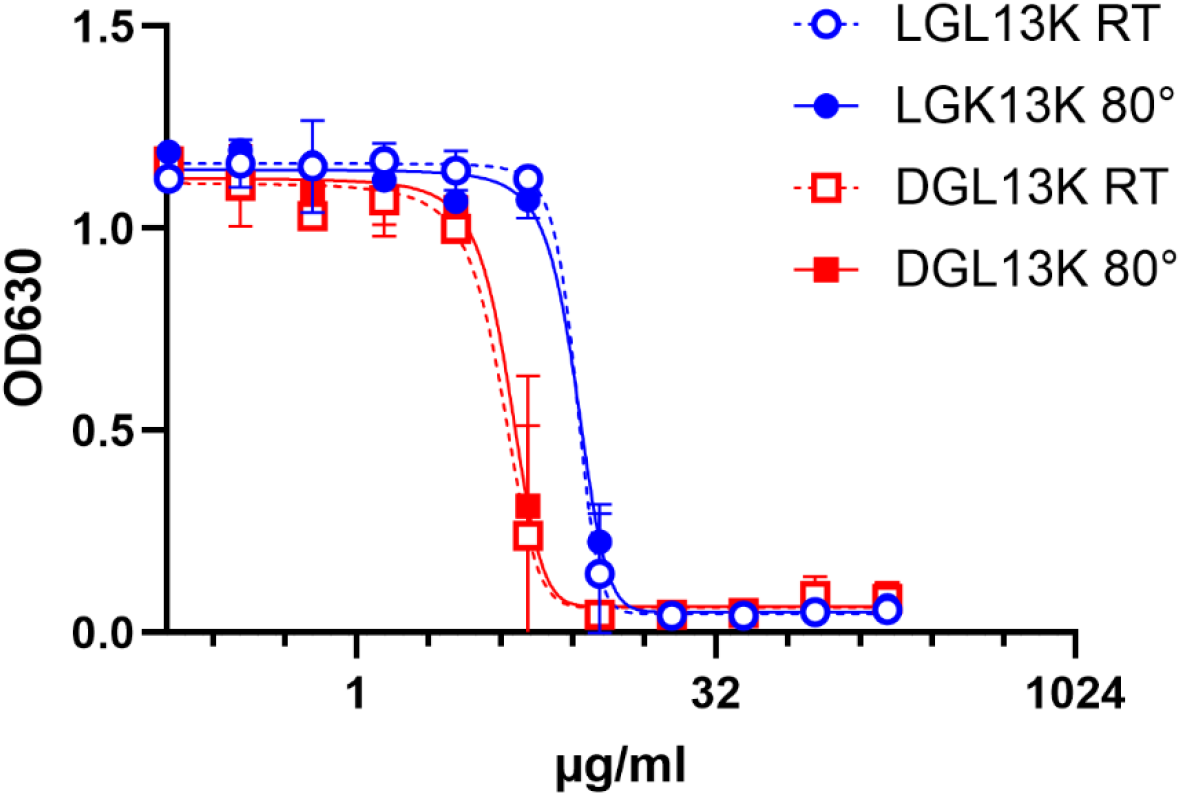
Heat stability of GL13K peptides. LGL13K (circles) or DGL13K (boxes) were incubated for 1 h at room temperature (open symbols) or 80°C (closed symbols) and then used for an MIC assay against *P. aeruginosa*. Duplicate samples were tested and plotted as mean ± range, N=2. The experiment was repeated with similar results.

### Peptide stereo chemistry

Proteolytic processing of AMPs has been cited as a concern for their development for clinical use [58,59]. DGL13K was originally designed to overcome proteolytic processing of the L-enantiomer in cultures of *P. aeruginosa* [16]. Indeed, DGL13K is not degraded in cultures of Gram-negative *P. aeruginosa* [16] and also resists proteolytic degradation in cultures of Gram-positive *Enterococcus faecalis* [19].

In addition to its greater stability, DGL13K also exhibits different antibacterial properties from the L-enantiomer. LGL13K is mainly active against Gram-negative bacteria while DGL13K also shows strong activity against Gram-positive bacteria [19,48] (**Table 2**). This difference may be due to the preferential binding of DGL13K to peptidoglycan, a component of the Gram-positive cell wall [60]. In contrast, both LGL13K and DGL13K bind to lipopolysaccharide, a component of the Gram-negative cell envelope [60]. Thus, the stereochemistry of the amino acids affects not only the stability of the peptide but also directly affects bacterial selectivity.

**Table 2:** MIC of peptide stereo-isomers. DGL13K, LGL13K, D-allo-Ile GL13K and L-allo-Ile GL13K were tested in MIC assays against *P. aeruginosa* and *S. aureus*. The median MIC is listed for 1-3 independent experiments. Pairs of D- and L-peptides (Ile vs. allo-Ile) were compared for each bacterial species by unpaired Student’s t-test. None were significantly different.

| Peptide | *P. aeruginosa* | N | *S. aureus* | N |
| --- | --- | --- | --- | --- |
| DGL13K | 7.8 µg/ml | 4 | 2.6 µg/ml | 6 |
| D-allo-Ile GL13K | 5.2 µg/ml | 3 | 2.6 µg/ml | 5 |
| LGL13K | 10.4 µg/ml | 4 | 83.3 µg/ml | 6 |
| L-allo-Ile GL13K | 10.4 µg/ml | 1 | 83.3 µg/ml | 3 |

In addition to the chiral centers at the alpha-carbon, GL13K contains two isoleucine residues that exhibit a second chiral center in the side chain. To test if this chiral center affects peptide activity, LGL13K and DGL13K were synthesized with two allo-Ile residues. MIC assays showed that for DGL13K and LGL13K for each bacterial species, the activity of the allo-Ile peptides matched that of the unmodified peptides (**Table 2**).

These results suggest that only the alpha-carbon chiral center affects peptide activity.

### Amino acid substitutions

The antimicrobial activity and toxicity profile of AMPs can be optimized by targeted amino acid substitutions. The antimicrobial activity of AMPs is typically defined by cationic and hydrophobic amino acids that target and disrupt the negatively charged bacteria membrane, respectively [30]. In this context, tryptophan and arginine have been identified as enabling peptide–membrane interactions and antibacterial activity [26,61-64] and were the focus of the tested modifications to DGL13K.

The two cationic amino acids lysine and arginine are often found in AMPs. DGL13K contains four Lys residues that contribute significant positive charge to the peptide (**Table 1**). In fact, without these Lys residues, the peptide does not exhibit bactericidal activity [34]. Lys can be substituted for the cationic amino acid Arg, which shows a linear relationship with hydrophobic residues in AMPs that does not exist for Lys [26]. Thus, Arg substitutions have been described to affect the antibacterial activity of AMPs [61,64].

Lys contains a four-carbon chain ending in a primary amine group with a pKa of 10.8 while Arg contains a 3-carbon aliphatic chain attached to a positively charged guanidinium group with a pKa of 12.5 [65]. To test if Arg substitution affected the activity of DGL13K, we designed DGL13R [66], which contains four Arg residues in place of the Lys residues in the parent peptide. The activity of DGL13R was not different from the parent peptide when tested against *S. aureus* (**Table 3**).

**Table 3.** MIC of modified peptide sequences. The MIC of DGL13K was compared to the Arg-substituted peptide DGL13R and the Trp-substituted peptide DGL12W. Each peptide was tested against *P. aeruginosa* or *S. aureus*. The median MIC is shown for 2-4 independent experiments. Values outside the tested range were set at 1000 µg/ml for calculation purposes. MIC values for *P. aeruginosa* were compared by unpaired Student’s t-test. *) different from DGL13K, P<0.01. n.d. - not determined. MIC values for *S. aureus* were compared to DGL13K by one-way ANOVA. **) different from DGL13K, P<0.0001.

| Peptide | *P. aeruginosa* | N | *S. aureus* | N |
| --- | --- | --- | --- | --- |
| DGL13K | 5.2 µg/ml | 4 | 1.3 µg/ml | 9 |
| DGL13R | n.d. |  | 2.6 µg/ml | 5 |
| DGL12W | 13* µg/ml | 6 | 917** µg/ml | 6 |

In addition to positively charged amino acids, hydrophobic amino acids play a role in membrane interaction. These can be grouped as aliphatic amino acids (e.g. Ala, Ile, Leu, Val) and aromatic amino acids (Trp, Phe). DGL13K contains seven of the former but none of the latter. Trp residues, in particular, have been introduced in AMPs [62,64] due to their preference for the interfacial region of lipid bilayers [61]. To examine the role of hydrophobic amino acids in peptide activity, DGL13K was redesigned by substituting one Ile and one Leu residue for Trp residues. To optimize the steric presentation of these amino acids, Lys residues were moved to generate a more amphipathic peptide [62] (**Table 1**). A helical wheel representation of the redesigned peptide, DGL12W, shows that charged and hydrophobic amino acids are now arranged on opposite sides of a predicted alpha-helix (**Fig. 2**).

**Figure 2:**
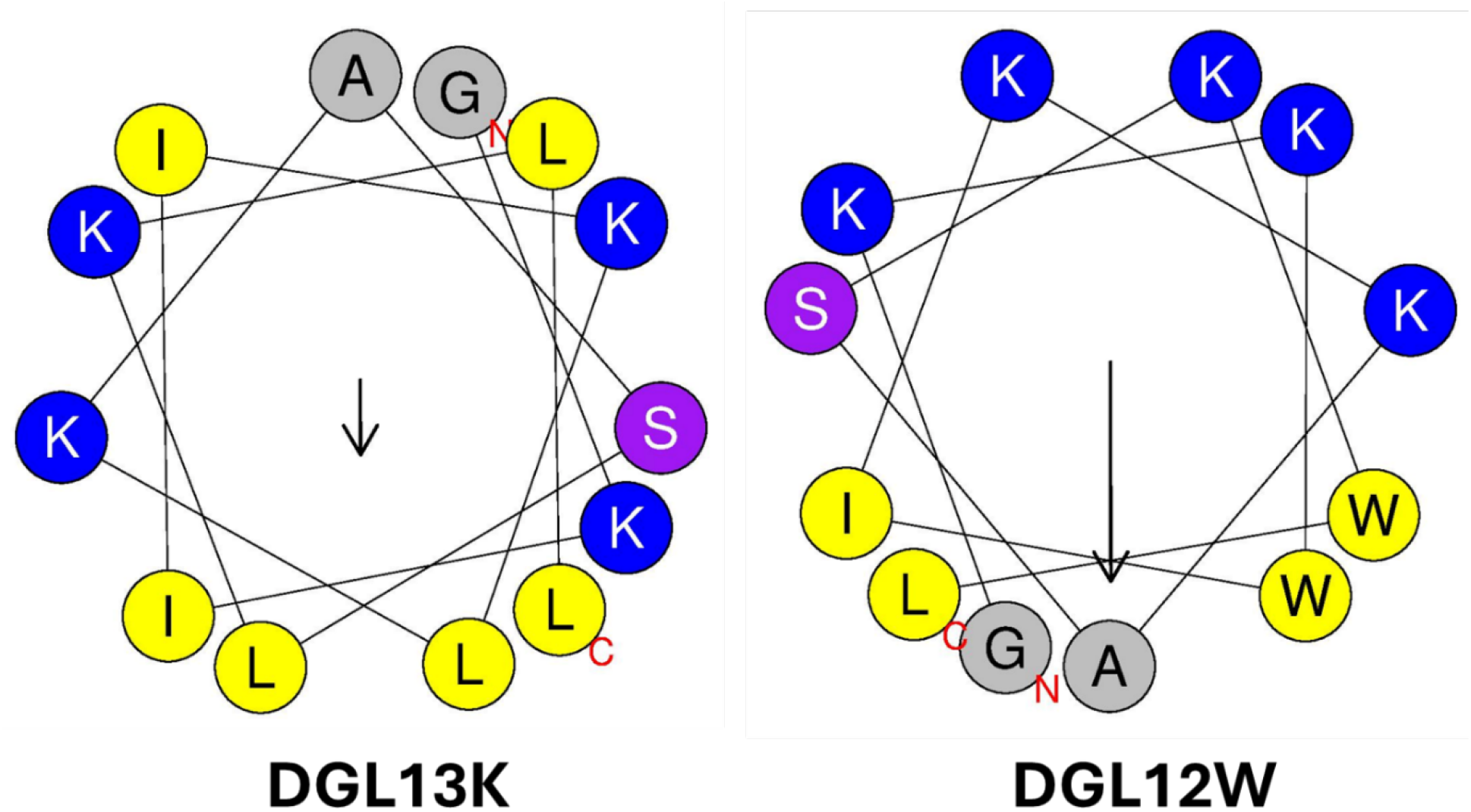
Helical wheel representations of DGL13K and DGL12W. Amino acid residues are labeled with the single letter code, the N- and C-terminal are labeled with red letters. Cationic amino acids are colored blue, hydrophobic amino acids are colored yellow and polar residues are purple. The images were produced using Heliquest [67].

DGL12W showed a 2-fold increase of the MIC against *P. aeruginosa* but had lost its activity against *S. aureus* with a mean MIC above the tested concentration range in some epxeriments (**Table 3**). Thus, re-arranging the location of the cationic residues and substituting two Ile/Leu residues with Trp created a peptide with increased specificity for the Gram-negative bacteria *P. aeruginosa*, compared to DGL13K. It is noted that these empirical changes are based on general rules for AMP design since no specific “AMP sequence motif” has been identified. As a result, each newly designed peptide sequence must be carefully tested to ensure that it exhibits the desired properties for stability, activity, specificity, toxicity and resistance.

### N-terminal modifications

The addition of functional groups, including polyethylene glycol (PEG) or myristoylate to the peptide sequence is known to affect peptide activity. For AMPs in particular, these modifications have been reported to increase antimicrobial activity and reduce peptide toxicity to mammalian cells [68-71]. To test the effect of these modifications on peptide activity, an N-terminally PEGylated version of LGL13K and N-terminally myristoylated and biotinylated (control) versions of DGL13K were prepared (**Table 1**) (**Fig. 3**)

**Figure 3:**
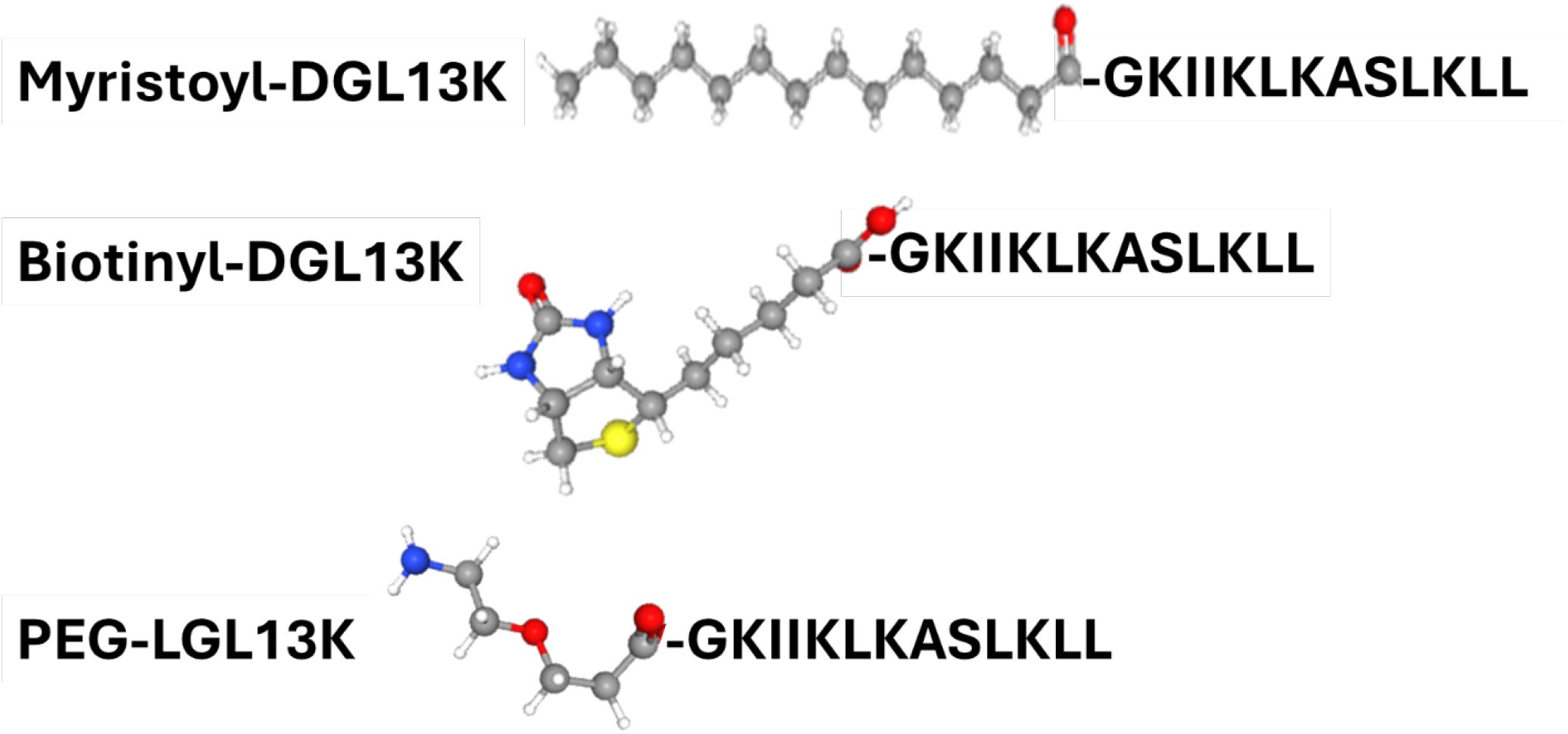
Structure of N-terminal modifications are shown in ball and stick models. Atom color code: Carbon - grey; hydrogen - white; oxygen - red, nitrogen - blue and sulfur - yellow. Peptide sequence is shown in one-letter code. Structures obtained from PubChem (https://pubchem.ncbi.nlm.nih.gov/)

The N-terminal PEGylation of LGL13K caused a 2- and 3-fold increase in MIC for *S. aureus* and *P. aeruginosa*, respectively (**Fig. 4**). Similarly, biotinylation increased the MIC for *S. aurues* and P. aeruginosa, 2- and 4-fold, respectively. In contrast, the addition of myristoylate to the N-terminus of DGL13K abolished its activity against both *P. aeruginosa* and *S. aureus*. Since the biotin molecule resembles the structure of PEG rather than that of myristate, these results suggest that addition of a highly hydrophobic chain inactivates the antimicrobial activity of the peptide sequence whereas addition of more polar molecules, which include oxygen and NH groups has only a minimal effect on peptide activity.

**Figure 4:**
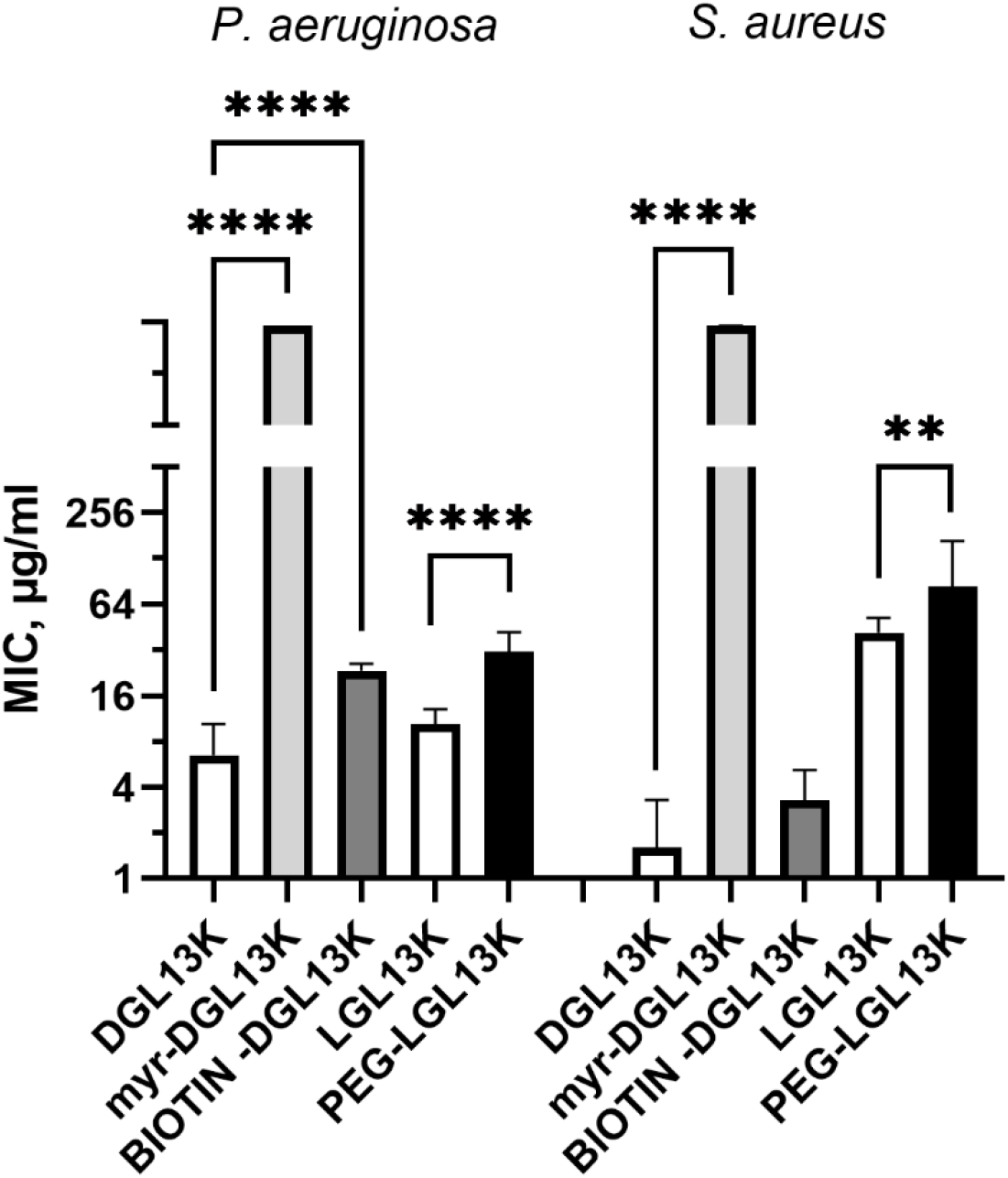
MIC of N-terminally modified DGL13K. The MIC against *P. aeruginosa* and *S. aureus* were determined for myr-DGL13K, biotin-DGL13K and compared to DGL13K in 2-6 independent experiments, which were analyzed by one-way ANOVA. PEG-LGL13K was compared to LGL13K in 3-5 independent experiments, which were analyzed by unpaired Students t-test. MIC values outside of the measured range were set at 1000 µg/ml for calculation purposes. ****) P<0.0001; **) P=0.005.

### Hemolysis

The selective activity of many AMPs exploits the compositional differences between prokaryotic and eukaryotic membranes to avoid mammalian cell toxicity. One measure of this toxicity is the peptide concentration leading to 50% lysis of red blood cells (HC50 = hemolytic concentration 50) [72]. Dose response experiments with DGL13K and LGL13K had revealed that HC50 is 500-1000 µg/ml [48]. To compare multiple peptides, each peptide was incubated with human red blood cells at 500 µg/ml and hemolysis determined spectrophotometrically (**Fig. 5**). DGL13K and LGL13K showed similar lysis as previously reported, while the Arg-modified peptide DGL13R exhibited a small increase in lysis. The rearranged peptide DGL12W, on the other hand, caused less than 10% hemolysis at this concentration. Thus, the higher bacterial selectivity of this peptide also resulted in improved selectivity for bacterial membranes, i.e. less erythrocyte toxicity. The PEGylated LGL13K peptide appeared to show less hemolysis than the unmodified peptide but this difference did not reach statistical significance.

**Figure 5.**
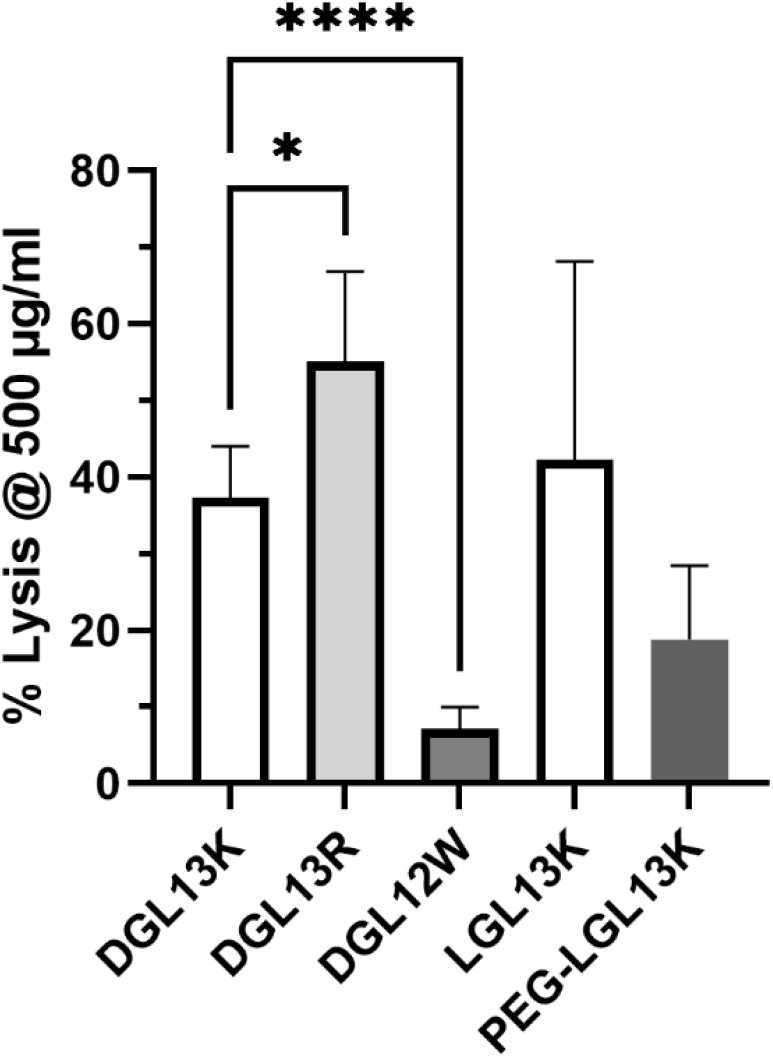
Hemolysis. The ability of different peptides to lyse human red blood cells at 500 µg/ml. The data from 1-6 independent experiments were plotted as mean ± 95% confidence interval and analyzed by one-way ANOVA for modified DGL13K peptides compared to unmodified DGL13K (open bar) and unpaired Student’s t-test for LGL13K peptides. *) P<0.02; ***) P<0.0001; N=2-13. A statistical outlier was removed using the ROUT method, Q-1% (Graphpad Prism 10.4).

It is noted that the hemolytic concentration used here is about 100-fold higher than the MICs determined for several of these peptides (**Table 2-3**), suggesting a high safety margin for clinical application. Indeed, we have found that topical application of 1 mg/ml DGL13K does not cause acute skin toxicity [48].

### Serum activity

The relatively high safety margin for hemolysis, suggested that the peptides could have systemic applications. To explore this option, antibacterial activity of select peptides was compared in the presence and absence of 50% serum. **Fig. 6** shows that the antibacterial activity against *P. aeruginosa* is lost in the presence of 50% serum, suggesting that the interaction with the Gram-negative cell envelope is highly sensitive to serum components. We have previously formulated DGL13K with EDTA to enhance antibacterial activity against *P. aeruginosa* [48], suggesting that divalent cations, e.g. calcium, could play a role in this interaction. Interestingly, the effect of serum on the antibacterial activity was more modest for *S. aureus* (**Fig. 6**). Several peptides, which showed low initial activity, were not affected by the presence of serum in the assay. These results support the view that the stereo-specific interactions of LGL13K and DGL13K with components of the Gram-negative and Gram-positive cell envelopes [60] are also differentially affected by serum.

**Figure 6:**
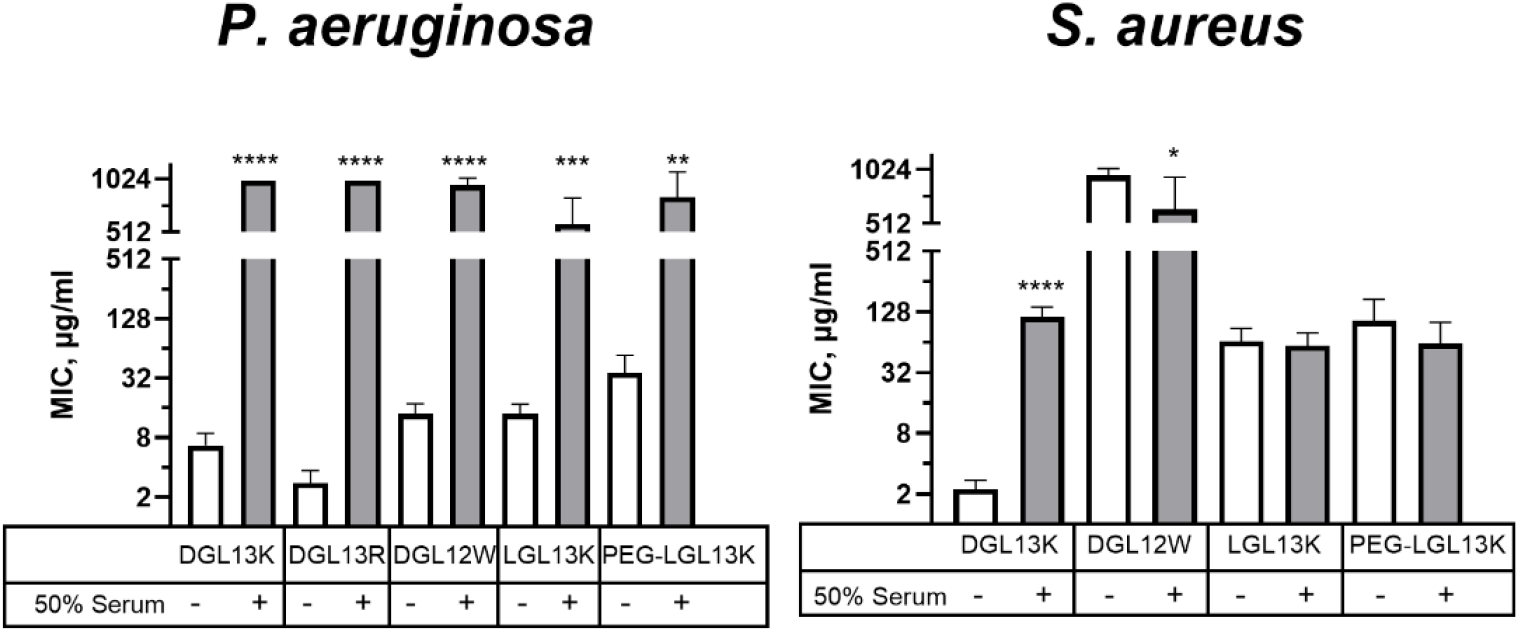
Relative activity of peptides in 50% serum. MIC of selected peptides were determined in the absence (open bars) or presence (shaded bars) of 50% serum for *P. aeruginosa* or *S. aureus*. Each sample [pair was analyzed by unpaired Studnets t-test with Welch’s correction for different variances, as needed. MIC values outside of the measured range were set at 1000 µg/ml for calculation purposes. *P. aeruginosa*: data from 1-9 independent experiments are plotted as mean ± 95% confidence interval. **) P<0.002; ***) P=0.0002; ****P<0.0001 (N=2-18). *S. aureus*: data from 2-10 independent experiments are plotted as mean ± 95% confidence interval. *) P<0.05; ****P<0.0001 (N=4-21).

## Conclusions

The original design of the GL13 peptide family was based on the location of a potential anti-inflammatory peptide in the sequence of the salivary protein BPIFA2 [46]. Thus, peptide GL13NH2 exhibited anti-inflammatory activity that captures the activity of the parent protein since GL13NH2 blocks the binding of LPS to BPIFA2 [44]. The substitution of three polar amino acids in GL13NH2 for cationic amino acids (Lys/Arg) results in an antibacterial peptide, LGL13K (Formerly GL13K, [34]. The present study demonstrates that this sequence exhibits optimized properties, presumably derived from the natural evolution of the parent protein sequence. Thus, none of the introduced modification were able to substantially alter the biological properties of GL13K. The modified peptides showed no more than a 2-fold increase of antibacterial activity, reduction of hemolytic activity or activity in the presence of serum. The specific interaction of each peptide enantiomer with the cell envelopes of Gram-negative and Gram-positive bacteria, deserve additional exploration as it may lead to the design of optimized AMPs with improved activity in serum [64].

The low serum activity is not unique to DGL13K and may be an inherent evolutionary aspect of these peptides, which are typically found in mucosal and skin surfaces [73]. Surprisingly the cationic peptide LL-37, which is found in circulating immune cells, also displays poor activity in the presence of serum; reviewed in [74]. Thus, it should be considered that in vitro assays with serum may not be a good approximation for in vivo activity in circulation.

Despite the low activity in the presence of serum, DGL13K shows activity in wound infections [48] and retains activity in synovial fluid (Gorr, unpublished) while LGL13K inactivates LPS in the peritoneum [34]. Given the stability and lack of bacterial resistance, in vivo activity and low in vivo toxicity of DGL13K, this peptide appears to be naturally optimized for the treatment of topical and localized infections.

## AUTHOR CONTRIBUTIONS

SUG is solely responsible for the content of this manuscript

## FUNDING

This study was supported with research funds from the University of Minnesota School of Dentistry.

## CONFLICT OF INTEREST

SUG is Chief Scientific Officer of and holds equity in Gavia BIO, LLC, which is developing the antimicrobial peptide Minneganan (DGL13K). These interests have been reviewed and managed by the University of Minnesota in accordance with its Conflict of Interest policies.

